# Ecological homogenisation in North American urban yards: vegetation diversity, composition, and structure

**DOI:** 10.1101/061937

**Authors:** William D. Pearse, Jeannine Cavender-Bares, Sarah E. Hobbie, Meghan Avolio, Neil Bettez, Rinku Roy Chowdhury, Peter M. Groffman, Morgan Grove, Sharon J. Hall, James B. Heffernan, Jennifer Learned, Christopher Neill, Kristen C. Nelson, Diane E. Pataki, Benjamin L. Ruddell, Meredith E. Steele, Tara L. E. Trammell

**Author notes:** Contributed equally to this manuscript.

## Abstract

Urban ecosystems are widely hypothesised to be more ecologically homogeneous than natural ecosystems. We argue that urban plant communities assemble from a complex mix of horticultural and regional species pools and thus evaluate the homogenisation hypothesis by comparing cultivated and spontaneously occurring urban vegetation to natural area vegetation across seven major US cities. Urban yards were homogenised across cities in terms of their diversity, composition, and structure. First, cultivated and spontaneous yard flora had higher numbers of species than did natural areas but similar phylogenetic diversity, indicating that yard species were drawn from a relatively small number of lineages. Second, yards were compositionally more similar across regions than were natural areas. Finally, vegetation structure, specifically cultivated tree density, was less variable in yards than natural areas across cities. Biodiversity homogenisation likely reflects similar horticultural source pools, homeowner preferences, management practices, and environmental filters across US cities.

## Introduction

The majority of humans now live in urban environments, and both urban area and population size are projected to increase (UN 2014), creating a pressing need to understand ecological processes within cities. Despite increasing urbanisation, its potential impacts on community assembly of organisms, biodiversity and ecosystem function are unclear—urban ecosystems were the least-studied in a recent review of over 11,500 assemblages (Newbold *et al.* 2015). Compounding this uncertainty, any urban flora is comprised of both human-cultivated and spontaneously occurring (establishing without human assistance) species, each of which is subject to distinct ecological and human influences (Knapp *et al.* 2012). While there is evidence of high biodiversity within cities (McKinney 2006; Grimm *et al.* 2008; Knapp *et al.* 2008; Newbold *et al.* 2015), few studies distinguish among these different aspects and drivers of urban biodiversity. This knowledge gap makes it difficult to interpret comparisons of diversity, both among different urban assemblages and when comparing urban and natural assemblages. There is, therefore, a need to better understand the structure and composition of urban ecosystems by considering the different vegetation pools (cultivated and spontaneous) present in urban areas.

Urban plant communities are a consequence of the ecological assembly processes that operate in natural ecosystems [*e.g.*, habitat filtering—Kraft *et al.* (2015) and competition— HilleRisLambers *et al.* (2012)], as well as human desires and influences. In Figure 1 we outline a conceptual framework describing how various filters act on the natural continental and horticultural industry plant species pools that together constitute the source pool for the assembly of urban plant communities. We focus on residential landscapes (*i.e.*, yards), under the assumption that residents have the greatest agency over, and frequency of interactions with, their own yards, making yards one of the front-lines of urban change. We focus solely on the species pools from which communities are assembled, and do not address local processes that may be operating within communities.

The major sources of urban flora we outline in Figure 1 are subject to contrasting filtering processes, each of which varies with spatial scale and likely by geographic region. The horticultural flora is influenced by accessibility of plant material, propagation constraints, and human preferences, and is further filtered by regulation and management processes. In contrast, the naturally assembled continental and regional floras are influenced by historical biogeographic processes and filtered by dispersal limits, climate, pollution, soil, and other abiotic constraints (Weiher & Keddy 2001; Ricklefs 2004; Wiens *et al.* 2010; Liu *et al.* 2011). Similarly, cultivated and spontaneous pools within the regional and urban flora are also subject to contrasting dispersal and filtering processes within the urban environment. For example, cultivated species are likely filtered by human preferences and management, and often receive additional resources (*e.g.*, water and fertiliser). Spontaneously regenerating species are likely also filtered by human management (*e.g.*, mowing, fertilising, and irrigating) and broader urban environmental conditions (*e.g.*, pollution and the urban heat island effect; Arnfield 2003). However, cultivated species may ‘escape’ cultivation to become part of the spontaneous urban and surrounding natural areas, becoming part of the wider regional species pool (Mack & Lonsdale 2001; Knapp *et al.* 2008). Natural areas surrounding cities are thus mixed assemblages derived from these separate species pools and the interactions between them.

Urban areas are frequently described as *homogenised* (ecologically similar; McKinney 2006; Grimm *et al.* 2008; Groffman *et al.* 2014), but it is unclear which of the many components of biodiversity (Purvis & Hector 2000) are subject to homogenising pressures, what factors contribute to such pressures, and how we might recognise patterns of homogenisation. In Box 1, we consider three potential aspects of urban vegetation that might exhibit homogenisation—*diversity, composition*, and *structure*—and present hypotheses regarding how urbanisation might influence these different aspects for both the cultivated and spontaneous vegetation pools. In describing homogenisation, we focus on urban cultivated and spontaneous species pools in relation to natural area pools. Homogenisation might be seen as a reduction in the number of lineages represented, more similar species compositions, or lesser variation across urban areas in structural aspects of the vegetation, including the overall height of vegetation. Contrasting urban and natural vegetation pools is key to our framework: the extent of similarity among natural systems reflects natural climatic, ecological and biogeographic processes, and it is critical to test whether urban systems show greater similarity than expected given these factors. Natural assemblages change along both micro-and macro-scale environmental gradients (Levin 1992; Chave 2013); testing whether urban systems show similar trends therefore requires comparable surveys of the surrounding vegetation.

Here we present results from a survey of urban vegetation diversity, composition, and structure in residential parcels (“yards”) in seven major US cities (Boston, Baltimore, Los Angeles, Miami, Minneapolis-St Paul, Salt Lake City, and Phoenix). Our survey covered broad environmental gradients and included comparable natural reference sites, permitting us to compare natural area vegetation and spontaneous and cultivated species pools in urban yards. By empirically evaluating our framework, we hope to shed light on the human contributions to ecological assembly processes in urban systems that influence biodiversity and ecosystem function.

## Box: Hypothesised urban vegetation structure and homogenisation among species pools

Understanding ecological assembly processes in urban systems and how they differ from surrounding natural areas requires consideration of the different filters and human factors that affect cultivated and spontaneous vegetation (Figure 1). Below, we examine three components of urban vegetation—*diversity* (species richness and phylogenetic diversity), *composition*, and *structure*—and describe how each might vary across the species pools we outline in our framework (Figure 1). We compare urban spontaneous and cultivated vegetation to natural area vegetation across a broad gradient in aridity, a major environmental gradient known to strongly influence naturally assembled plant communities.

Understanding urban vegetation in the context of surrounding natural areas requires an understanding of the different filters and processes that affect the assembly of cultivated and spontaneous species (Figure 1). Below, we examine three components of urban vegetation— *diversity, composition,* and *structure*—and describe how each might vary across the species pools we outline in our framework (Figure 1). We focus on vegetation as a function of water stress (aridity), as this is the major environmental gradient we test empirically across our seven major metropolitan areas.

### Diversity

Human transport and management (*e.g.*, irrigation) of vegetation enables cultivated species to overcome natural dispersal and establishment barriers, such that we expect the species richness of the cultivated urban pool to be higher than that of urban spontaneous or nearby natural pools. Plant species richness of all species pools should positively correlate with moisture availability, consistent with well-established relationships between species richness and climate stress (*e.g.*, Currie 1991; Wiens & Donoghue 2004; Fine 2015). If humans prefer variation and can irrigate to overcome water limitation, we might expect the cultivation of a wide diversity of plant lineages to increase phylogenetic diversity in cultivated pools. If irrigation cannot overcome water limitation, we would expect the natural and cultivated pools’ phylogenetic diversity to mirror each other. In such a case, *in situ* diversification and historic biogeography would interact to determine phylogenetic diversity (Webb *et al.* 2002; Cavender-Bares *et al.* 2009). Were a limited subset of the tree of life cultivated in comparison with the natural pools, this would represent a form of phylogenetic homogenisation.

### Composition

If climate is a strong filter on the composition of spontaneous and natural area species pools, we expect the species and phylogenetic clades (the composition) of pools to vary across regions. However, within regions, these pools are drawn from the regional flora and are subject to the same (or similar) climate filters, so we expect some compositional similarity within regions. By contrast, if human preferences, transport, or management (*e.g.*, irrigation) relax the constraints imposed by climate and dispersal barriers, we expect cultivated pools to be homogenised: more similar to one another among regions than the spontaneous or natural area pools. We expect the spontaneous pool to be intermediate in composition to the cultivated and natural pools if it receives propagules from both pools and/or facilitates dispersal between the cultivated pool and the natural areas pool.

### Structure

Given similar human preferences (*e.g.*, savanna-like yards) and management (*e.g.*, irrigation) to mitigate climatic constraints, we expect cultivated pools to have similar structures (*e.g.*, tree height and density) across regions. By contrast, we suggest climate filters will lead to divergence in the structure of natural area pools, with taller trees, and greater tree density in wetter compared to more arid regions. Spontaneous pools could be intermediate between natural areas and cultivated pools as they are subject to less management than cultivated pools, but more management than natural areas. Lower variance in structural attributes (greater structural similarity) among urban pools compared to natural pools would suggest less variation in urban vegetation structure, and represent a form of homogenisation.

**Figure 1:**
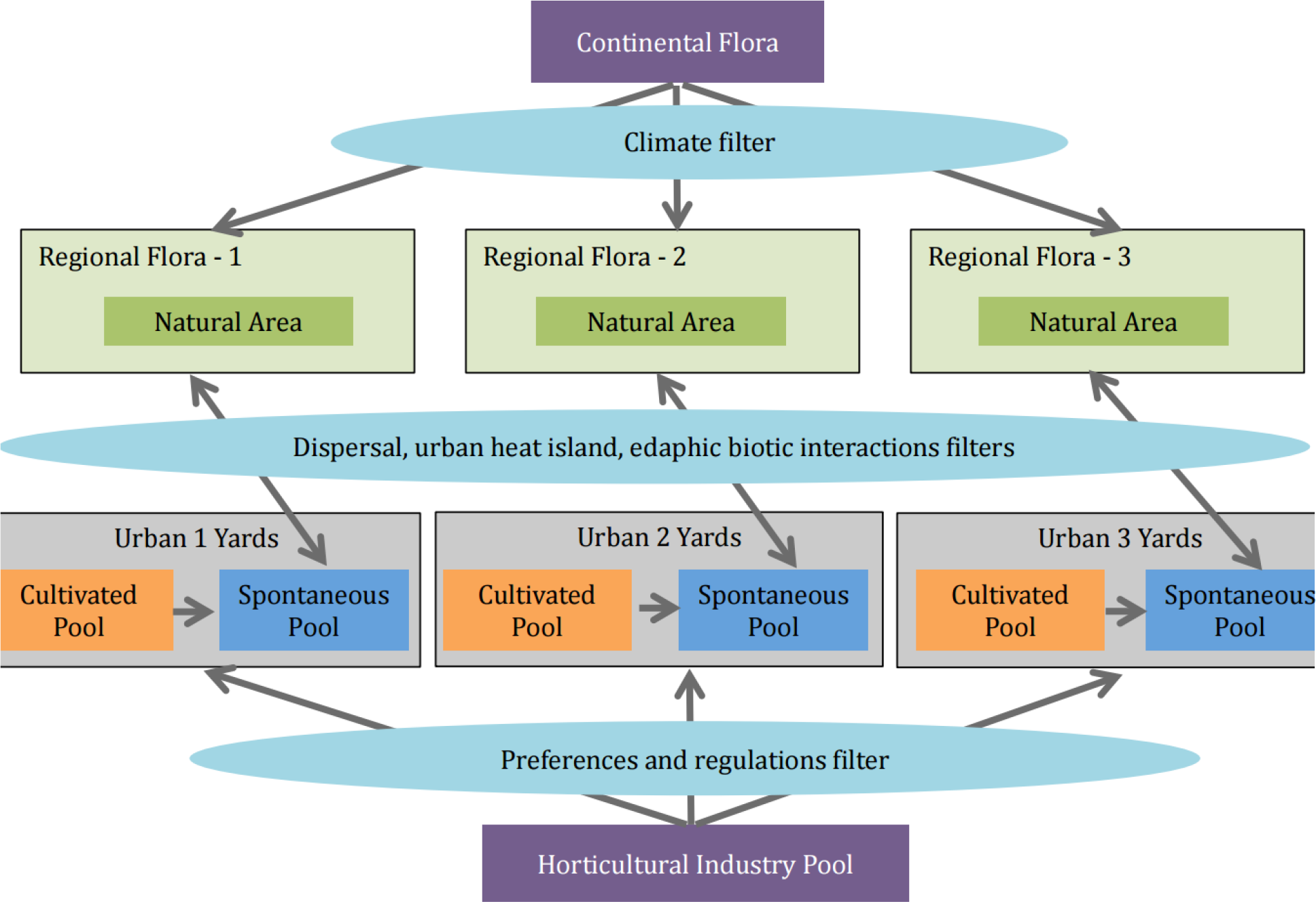
Conceptual overview of urban community assembly. Community assembly of the urban yard flora is driven by the movement of plants from the regional flora and the horticultural industry pool through contrasting filters. The regional flora are subsets of the continental flora, filtered by climate and dispersal limitations, and include natural vegetation pools that have emerged through a combination of dispersal and *in situ* speciation. The urban spontaneous flora assembles through dispersal from the regional flora, excluding species that cannot disperse into or persist in the urban environment given the urban abiotic and biotic pressures. The spontaneous flora also includes species that have escaped human cultivation and are able to propagate and establish on their own. The cultivated urban flora are largely a subset of the species available from the horticultural industry, filtered by human preferences and policy regulations that limit planting of invasive species. Species from the horticultural pool can assemble and become established in the regional and continental flora through migration into and out of the urban spontaneous pool.

## Methods

The sampling design of the Urban Homogenization of America project has already been described (Groffman *et al.* 2014; Polsky *et al.* 2014), but we outline it briefly below. Within each of seven major US metropolitan areas (Boston, Baltimore, Los Angeles, Miami, Minneapolis-St Paul, Salt Lake City, and Phoenix) we identified 21-30 urban household yards and 3-6 natural area sites. Within each metropolitan area we collected the vegetation (species presence/absence), tree structural trait, and leaf functional trait datasets that we describe in more detail below. All software described below are *R* (*v3.2.2*; R Core Team 2015) packages unless otherwise stated.

### Vegetation (species presence/absence) data

An exhaustive presence/absence survey was conducted in the yard of each household. For parcels with yards larger than 1.0 ha, yard components (front lawn, back lawn, woodland, woodlot or unmanaged, perennial bed, vegetable garden, xeriscape) were inventoried (but not analysed) separately. The entire area of each yard was surveyed except where there was an unmanaged vegetation or woodland/woodlot component, which was sampled via full-length or 100 m x 2 m transects, whichever was shorter. Species were designated as spontaneous or cultivated; a given species could be documented as both spontaneous and cultivated if different individuals of that species fell into different categories. Land-use and land-use history were considered in the designation. For example, species in woodlots and unmanaged vegetation components were generally considered spontaneous.

Between three and six natural areas were designated in each region, chosen to represent similar ecological regions and topographic and edaphic features of the urban region. Within each natural area, eight transects were established, each treated as a separate sample, for a total of 24-48 transects (100 m x 2 m), comparable to the household sample size. All vegetation in view from within the transect area was exhaustively recorded for species presence/absence. The locations and directions of the transects within the reference areas were randomly assigned in advance using GIS mapping. We caution that, while best efforts were made to select natural sites representative of vegetation before urbanisation, there are few (if any; Mann 2005) parts of North America not impacted by humans.

Species names were matched to The Plant List (http://www.theplantlist.org) version 1.1, using *Taxonstand* (Cayuela *et al.* 2012). The Zanne *et al.* (2014) phylogeny was used for all phylogenetic metrics, and species missing from this tree were bound in at the genus level using *pez*’s *congeneric.merge* (Pearse *et al.* 2015). Hybrids and species for which there were no phylogenetic data were excluded from the analyses.

### Tree structural trait data

For trees, data for number of individuals, diameter at breast height (DBH), height, and crown projected area for all trees in yards less than 0.1 ha were collected, following protocols developed by the US Forest Service for use with the UFORE models in their ‘iTree’ application (although we do not present iTree output here; http://www.itreetools.org). For large yards > 0.1 ha, 8 m radius plots were randomly established using GIS mapping at the ratio of 5 per hectare, rounded down to the nearest whole number. In natural sites, three 8-m radius plots were established per reference site for a total of nine to eighteen plots per region. No tree structural trait data were collected in Salt Lake City.

### Leaf functional trait data

Leaves were collected from three to five individuals per species, from three to five different households per city, whenever possible. One to three leaves were collected per individual, depending on size of plant. Sun leaves were collected, if possible. Heights and life form of each donor plant were recorded. All new species encountered in the reference sites were sampled; leaves were collected for three to five individuals per species across all reference sites, if possible. Leaves from a single individual were placed in a coin envelope placed on cardboard spacers fastened with rubber bands to press the leaves flat and absorb moisture prior to shipping to the University of Minnesota. No leaf functional trait data were collected in Salt Lake City; samples were collected in Phoenix, but were damaged in transit and so could not be analysed.

The leaf specimens were processed using a Python script (‘*stalkless*; (http://willpearse.github.io/stalkless). The *R* code in *stalkless* makes heavy use of *Momocs* (Bonhomme *et al.* 2014), which should be cited whenever it is used. Briefly, *stalkless* segmented the individual leaves present within each scanned image, identifying darker areas as objects (in this case leaves) with reference to the mean intensity of the image plus a multiple of the standard deviation of the image’s intensity (by default 2). Using *R*, candidate leaf images containing too much background noise or other objects in the scanner were removed by checking the dimensions of the images. A preliminary Fourier analysis using *eFourier* (Bonhomme *et al.* 2014) isolated remaining non-leaf images, grouping them together in a hierarchical cluster analysis of a Euclidean distance matrix of the Fourier parameters. We manually checked, verified, and supervised this process, which all *stalkless* users are strongly encouraged to do. These steps left us with images of 754 species (out of a total 2224 in the dataset) for the final analysis. We then used *stalkless* to record individual leaf surface area, perimeter length, and leaf compactness 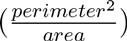. Smaller leaf surface area is associated with arid environments because of the reduced evaporation and water loss associated with small leaves. A higher perimeter per area would be expected in warmer climates because it is associated with either smaller leaves or more lobed leaves, both of which reduce boundary layer resistance and allow more rapid leaf cooling (Sack *et al.* 2003).

### Statistical analysis

Below, we present our analysis following the three groupings in box 1: *diversity, composition*, and *structure*. For all analyses, we ignored abundances in sampled assemblages and treated a species as present in a pool if it was recorded at least once in an assemblage associated with that pool type. Our seven cities lie along a major aridity gradient throughout continental North America. To quantify this gradient, Palmer Drought Severity Index (PDSI) data were downloaded from a global database maintained by the Consultative Group for International Agriculture Research (Trabucco & Zomer 2009) at a resolution of 30 arc-seconds. PDSI is a composite temperature-precipitation index that assesses how dry a particular region is; negative values indicate drier conditions, larger positive values wetter conditions.

*Diversity*. Species richness and phylogenetic diversity (mean phylogenetic distance; MPD) were calculated for each pool. Since MPD can scale with species richness, we report its standard effect size here (*SES_MPD_* Kembel 2009) calculated used *picante* (Kembel *et al.* 2010). We regressed the diversity metrics against aridity in each city, using mixed effects models where the aridity gradient and habitat pool were fixed effects and region was a random effect (using *lme4* and *lmerTest*; Bates & Maechler 2010; Kuznetsova *et al.* 2016). To test for changes in the variation of the diversity metrics within pools, we used Levene’s tests as implemented in *car* (Fox & Weisberg 2011).

*Composition*. We calculated the Sorensøn’s index of species compositional difference (using *vegan*; Oksanen *et al.* 2013), and the Phylosorensøn’s metric of phylogenetic distance (also using *picante*; Bryant *et al.* 2008; Kembel *et al.* 2010) for all the parcels. We modelled the Sorensøn’s distances among all pools as a function of whether those pools were of the same vegetation type *(*e.g.*,* spontaneous vs. cultivated) or region (*e.g.*, Boston vs. Phoenix). We then performed an ordination analysis for visualisation purposes (constrained to two dimensions, also using *vegan*; Oksanen *et al.* 2013).

*Structure*. We analysed four structural metrics: tree height, tree density (total number of trees divided by vegetative parcel area), leaf surface area, and leaf perimeter:area (a log_10_-transformed ratio). We obtained medians of each of the structural attribute either across all parcels in which a particular vegetation pool was measured (for tree density and height), or across all individuals measured (for leaf surface area and perimeter:area). We then treated and analysed these pool-level aggregates in exactly the same way as we analysed the diversity metrics above. Urban trees are easy to spot, and any unwanted spontaneously regenerating trees are both easy to identify and remove before they become large. We therefore, for the purposes of our tree structural metrics, treat all trees in urban areas as cultivated (*i.e.*, none are spontaneous).

## Results

### Diversity

In support of our hypothesis (Box 1), species richness was greater in the cultivated pool than the spontaneous pool, and the spontaneous pool had higher species richness than the natural area pool (Figure [diversity]a). In partial support of our hypothesis, both the spontaneous and cultivated pools had higher species richness in the less arid regions, while that of the natural species pool remained constant across regions(Figure 2a). In contrast to our hypothesis, there was no evidence of homogenisation of species richness across regions: species richness was no less variable among cities for the cultivated and spontaneous species in yards than for the natural area species.

However, there was evidence of homogenisation of phylogenetic diversity in yards across regions. Despite higher species richness of cultivated and spontaneous species in yards than in natural areas, phylogenetic diversity (*SES_MPD_*) did not differ among vegetation pools (Figure 2c & Figure 2d), nor did it vary systematically across regions (Figure 2b). Thus species in both the cultivated and spontaneous pools appear to be drawn from relatively fewer lineages than species in the natural areas. Note that full statistical support for these trends is given in Supplement S1.

### Composition

In support of our hypotheses (Box 1), species and phylogenetic composition of the cultivated and spontaneous species in yards were more similar across regions than the natural area pools (Figure 3), evidence of compositional homogenisation in urban yards. This represents a form of biotic homogenisation. However, those species found in natural areas were also present in urban pools: natural pool compositions were nested within urban pools (*NODF* = 9.93, greater than all 1000 trial swap null permutations). This was not true of phylogenetic compositions (*NODF* = 1.34): lineages found in natural areas were not necessarily a subset of lineages found in yards.

### Structure

The ratio of leaf perimeter to surface area significantly decreased across the PDSI gradient (Figure 4). There was no significant interaction between vegetation pool and PDSI in overall perimeter:area, suggesting both cultivated and natural vegetation responded to the aridity gradient similarly. However, in contrast to our hypothesis (Box 1), no other structural metrics (tree height, density, and leaf surface area) varied across the PDSI gradient (Figure 4). Equally, there were no significant differences in variance in the structural metrics between habitat pools (Figure 5). However, in the natural surroundings of Salt Lake City and Los Angeles there were no trees whatsoever and there were very few trees in natural areas around Phoenix, while yards in these regions had tree densities greater than zero. Thus, yards are qualitatively more similar in tree density across regions than natural areas, evidence of homogenisation of vegetation structure. As discussed in the methods, we recorded no urban trees as spontaneous (see Figures 4 & 5).

**Figure 2:**
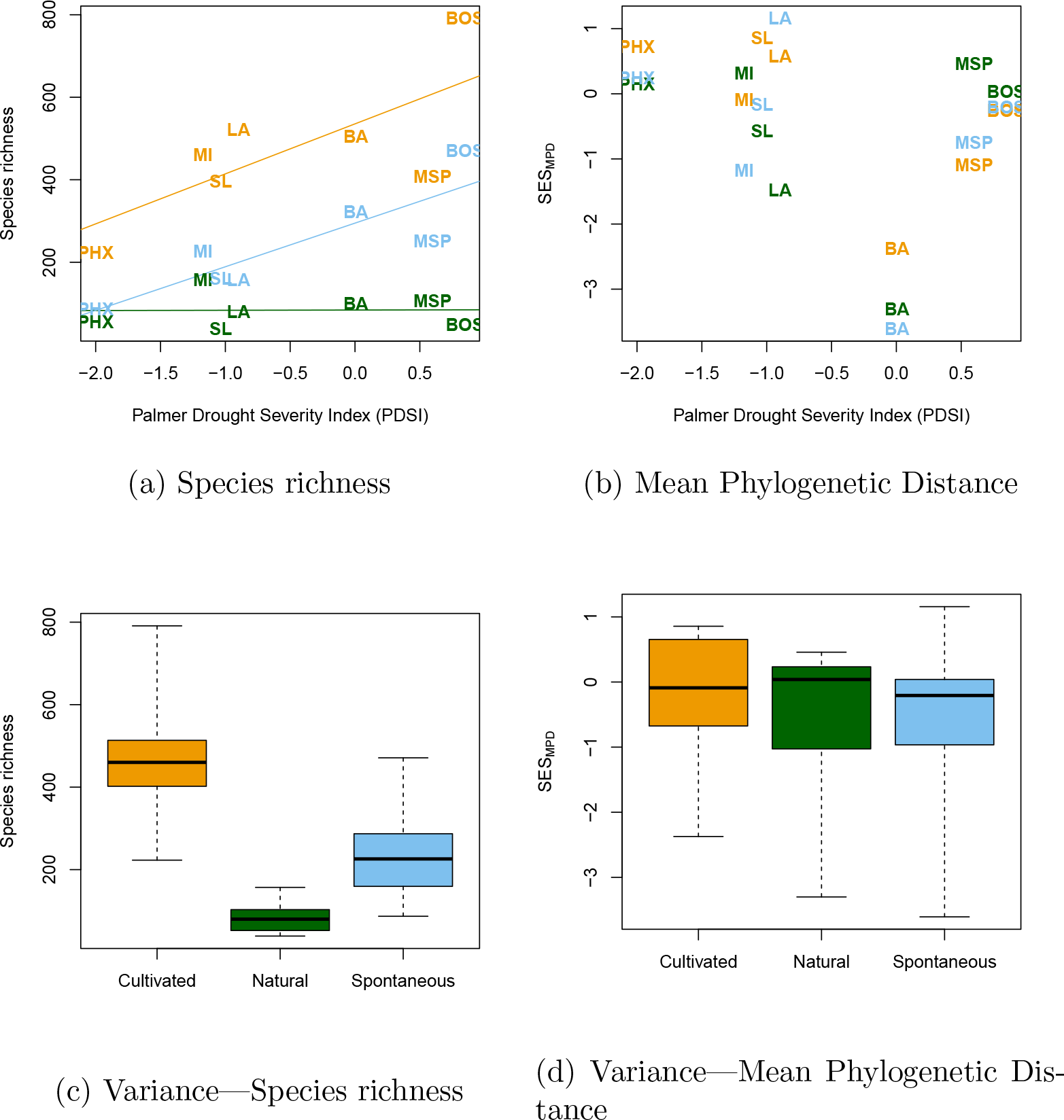
Diversity results. Regressions of species richness (a) and mean phylogenetic distance (MPD; b) against the aridity gradient. The three vegetation pools are represented by colour (cultivated species in yards, natural area species, and spontaneous species in yards in orange, green, and blue, respectively), and the regions themselves with abbreviations (Boston, Baltimore, Los Angeles, Miami, Minneapolis-St Paul, Phoenix, and Salt Lake City, as BOS, BA, LA, MI, MSP, PHX, and SL, respectively). There is support for significant differences in species richness among the three pools (*t*_9.986_ = 46.50, *p* ≤ 0.0001), and a significantly different response to the aridity gradient in the natural pool compared to the cultivated and spontaneous pools (*t*_9.986_ = −2.83, *p* = 0.012). There was no support for change in MPD across the aridity gradient (*t*_8.21_ = −1.30, *p* = 0.23) nor were there significant differences among vegetation pools (*t*_10.00_ = −0.58, *p* = 0.58). Full mixed effects model results are given in Supplement S1. Boxplots of the distributions of species richness (c) and MPD (d) in the three habitat pools (cultivated, natural, and spontaneous). The whiskers on the boxplots represent the limits of the data, and the boxes the inter-quartile range. There is no evidence for differences in variance across the vegetation pools in species richness (Levene’s test *F*_2,18_ = 1.71, *p* = 0.21) or MPD *F*_2,18_ = 0.03, *p* = 0.98). Apparent differences in variance in the figure are likely driven by non-Normality of the data, which the Levene’s test is not sensitive to.

**Figure 3:**
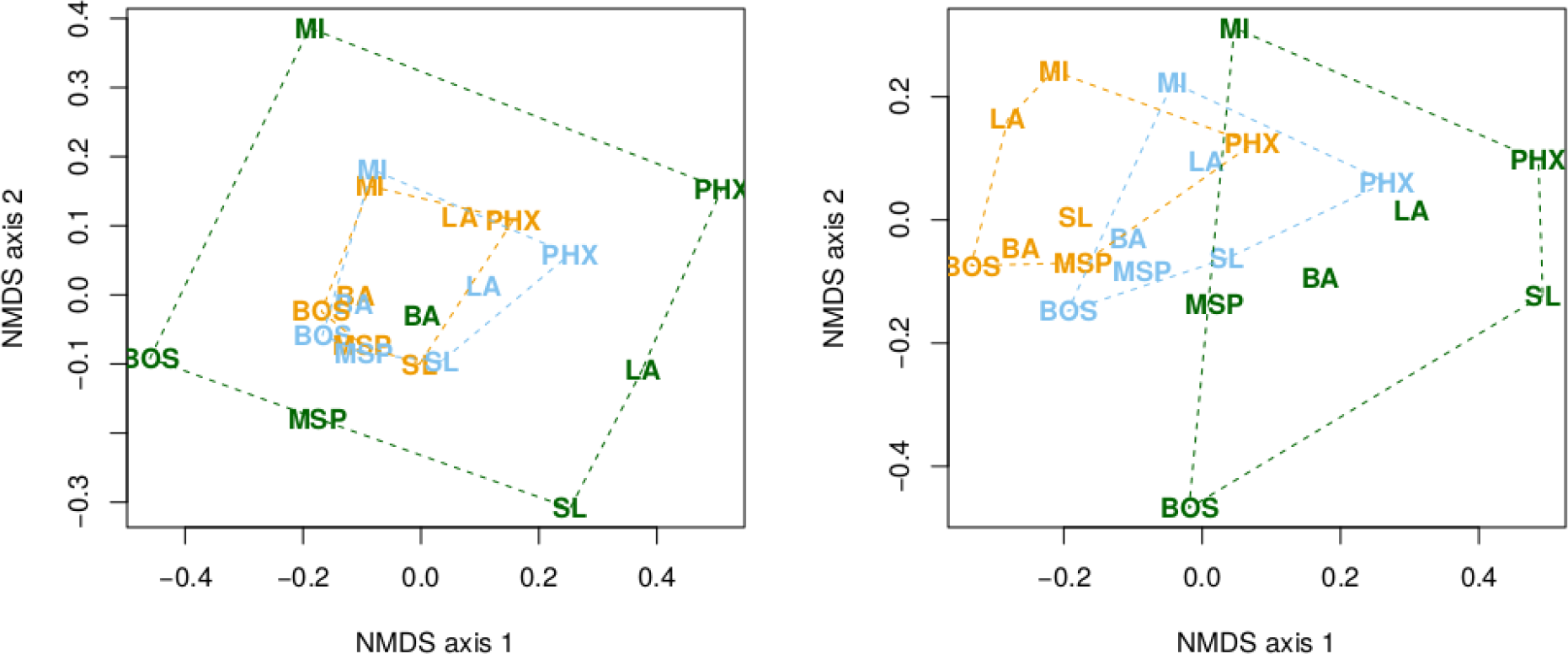
Composition results. Two axes of ordination of species (left; Sørenson’s distance) and phylogenetic (right; Phylosørenson’s distance) compositions across regions and vegetation pools. The three vegetation pools are represented by colour (cultivated, natural, and spontaneous in orange, green, and blue, respectively), and the regions themselves with abbreviations (Boston, Baltimore, Los Angeles, Miami, Minneapolis-St Paul, Phoenix, and Salt Lake City as BOS, BA, LA, MI, MSP, PHX, and SL, respectively). There is strong statistical support for an interaction between differences in vegetation pools and region under comparison in both species (*F*_11,141_ = 14.74, *r*^2^ = 0.54, *p* < 0.0001) and phylogenetic (*F*_11.141_ = 8.48, *r*^2^ = 0.0.40, *p* < 0.0001) structure. Cultivated and spontaneous pools are more similar across regions than natural area pools, and in all cases, pools in the same geographical area are more similar than pools across a geographical region (see Supplement S1). As discussed in the text, these figures are interpretative guides only; they are likely affected by biases in nestedness and artefacts from compressing dissimilarity into two dimensions for printing.

**Figure 4:**
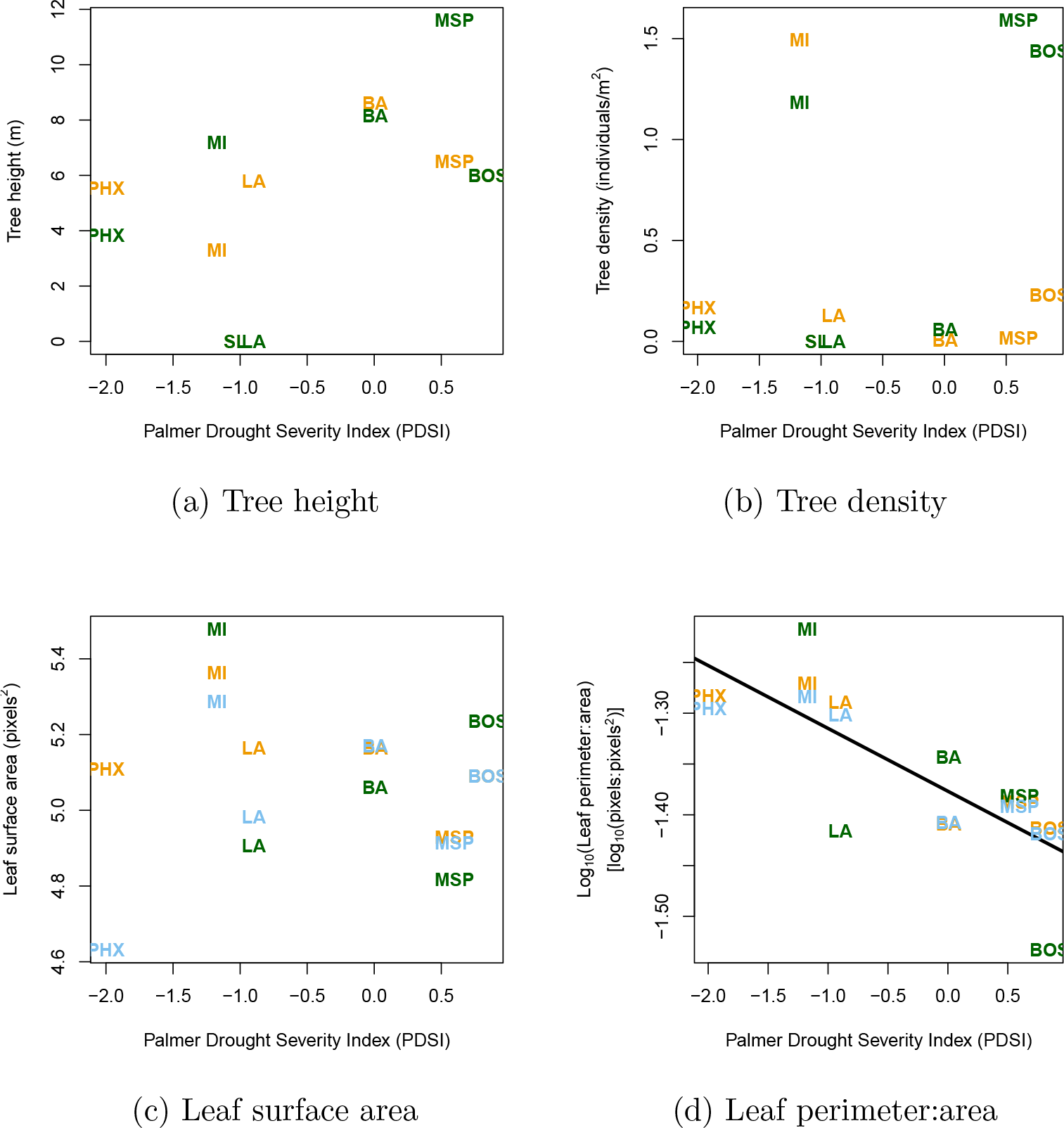
Structure results. Regressions of tree height, tree density, leaf surface area, and leaf perimeter:area ratio against the aridity gradient. The three vegetation pools are represented by colour (cultivated, natural, and spontaneous in orange, green, and blue, respectively), and the regions themselves with abbreviations (Boston, Baltimore, Los Angeles, Miami, Minneapolis-St Paul, Phoenix, and Salt Lake City, as BOS, BA, LA, MI, MSP, PHX, and SL, respectively). While *log*_10_(perimeter:area) significantly changed across the gradient (*t*_4.43_ = − 3.90, *p* = 0.015), there were no other statistically significant differences either across the aridity gradient or among habitat pools. Note that, in the text, we discuss that neither Los Angeles nor Salt Lake City had any trees in their natural pools. See Supplement S1 for full mixed effect models results.

**Figure 5:**
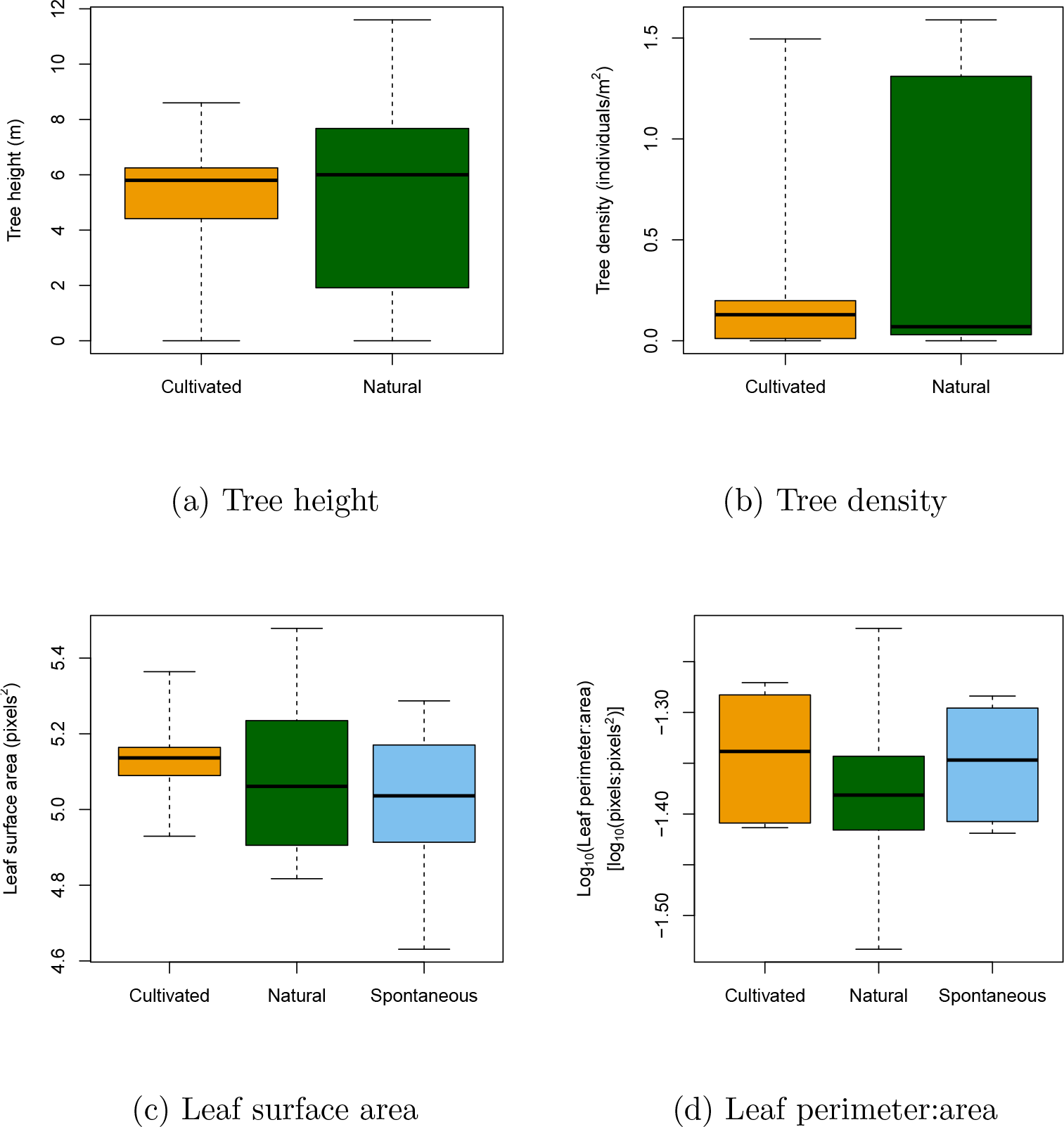
Structure—variance results. Boxplots of tree height, tree density, leaf surface area, and leaf perimeter:area ratio for the three vegetation pools (cultivated, natural, and spontaneous). The whiskers on the boxplots represent the limits of the data, and the boxes the inter-quartile range. Note that, as discussed in the main text, there are no tree structural metrics for the spontaneous pool. There is no statistical support for unequal variances in any of these variables (all Levene’s test *F*_1,12_ < 1.71, *p* > 0.20). We note that the full distributions of these data, and statistical analysis of differences in mean, are given in figure 4.

## Discussion

Urban plant assemblages are typically described as similar to one another, or ‘homogenised’ (McKinney 2006; Grimm *et al.* 2008; Groffman *et al.* 2014). Taking a species pool-based approach to plant diversity, we have confirmed that indeed, the composition of cultivated and, to a lesser degree, spontaneous urban plants is more similar among urban areas than among comparable natural reference areas. Furthermore, cultivated and spontaneous species in yards come from a more limited set of lineages (relative to their species richness) than do natural area species. Finally, tree density has a tendency to be more similar in yards than in natural areas across regions. However, despite management actions such as irrigation, aspects of plant structure in urban areas can vary across an aridity gradient, and, for some structural metrics, natural and urban plant pools are indistinguishable. Below we discuss these results, and argue that by distinguishing among plant diversity, structure, and composition, both variation and homogenisation can be detected within North American urban flora.

### Diversity

In keeping with other studies (reviewed in (reviewed in Pickett *et al.* 2001; Grimm *et al.* 2008), we found that urban vegetation, both in terms of the cultivated and spontaneous pools of species, had greater species richness than the natural areas, (Figure 2). However, phylogenetic diversity (*SES_MPD_*) showed no variation among vegetation pools. This suggests that, while cities may contain more species because of cultivation, these species come from a relatively limited set of lineages compared with species in natural areas. This (and other lines of evidence; Knapp *et al.* 2012) suggests that the concept of homogenisation with respect to the diversity of species in cities needs to be subtly refined. Urban residents cultivate many more species than occur in natural areas, but cultivated species comprise a smaller fraction of the tree of life than would be expected based on observations in natural vegetation.

Surprisingly, the species richness of the natural pools across the aridity gradient showed no relationship with the PDSI, while richness of the cultivated and spontaneous pools increased as aridity decreased (Figure 2). Despite homogenised lawn management and irrigation across the USA (Polsky *et al.* 2014), cultivated and spontaneous species richness responded to the aridity gradient (Figure 2). That we found change along the aridity gradient suggests that human management has not completely overcome the gradient. Multiple environmental gradients across North America drive patterns of species diversity (O’Brien *et al.* 2000); overcoming each of these gradients may require time, effort, and money that homeowners are unwilling to spend. In addition, nursery stock might be limited in more arid regions. Plant functional characteristics are known to display strong phylogenetic conservatism (Cornwell *et al.* 2014; Pennell *et al.* 2014); by sampling a relatively limited subset of the tree of life householders may make it harder to match species’ tolerances and environmental conditions.

### Composition

There is strong evidence that the cultivated and spontaneous species pools in yards are homogenised across regions relative to the natural area vegetation in terms of both species and phylogenetic composition (Figure 3), providing evidence for the claim that species within cities are similar (McKinney 2006; Grimm *et al.* 2008; Aronson *et al.* 2014). For the cultivated pools, this homogenisation might arise because plant nurseries offer a similar suite of species across the country, resulting in a homogenous source pool, or because of similar preferences across regions. For the spontaneous pool, homogenisation might result from similar filtering processes imposed by cities across regions, such as mowing, irrigation, and the urban heat island. Interestingly, the cultivated and spontaneous pools within the same region were similar to each other (see Figure 3), perhaps reflecting the influence of filtering by the extreme climate variation across regions and possibly the escape of cultivated species into the spontaneous pool within cities. More fundamentally, these results reflect the reality of the urban composition of North America: in part homogenised, in part regionally variable, as a consequence of both environmental filtering processes (driven by factors such as aridity) and human preferences.

The nestedness of regions’ species compositions (Figure 3) further complicates a simplistic pattern of homogenisation throughout North America. That natural areas should be nested within urban areas reveals that there are relatively few species present only in natural areas. We emphasise that this is a general trend in the composition of species pools across cities, and does not imply that natural assemblages, with their characteristic ecosystem functions, are found in urban areas. Rather, it implies a dis-assembly of natural ecosystems into *hybrid* ecosystems (Hobbs *et al.* 2009): ecosystems containing some exotic and some natural species. These data are dominated not by homogenisation, but *conurbation*, whereby the previously unique and independent natural habitat pools surrounding urban areas are merged and combined. Such mixing has profound implications for species’ future evolution, breaking down existing species-associations (essentially invasional meltdown; Simberloff & Holle 1999), and through increasing diffuse interactions makes the evolution of density-dependent competitive interactions and Janzen-Connell effects difficult (or impossible; Zillio *et al.* 2005; Hubbell 2008). Such hyper-diverse mixtures of species that have not previously interacted could have profound implications for surrounding natural regions, and long-term change within cities.

### Structure

The lack of statistically significant differences in the means and variances in tree density and height across natural and cultivated pools are, we argue, a product of quantitatively incomparable data. The three cities surrounded by desert (Phoenix, Salt Lake City and Los Angeles) all had tree densities at or very nearly zero trees per hectare in natural areas, whereas trees were common in yards in these regions. We suggest that the placement of trees in urban areas surrounded by desert is sufficient to argue for a homogenisation of tree density in our dataset. We also suggest our comparatively small sample size of cities (necessary given the scale of fieldwork required to survey major metropolitan areas) means we have reduced statistical power. Thus we argue that these results reflect homogenisation of urban vegetation on the basis of properties like tree cover that stakeholders perceive to regulate ecosystem services (*e.g.*, climate regulation and aesthetics; Larson *et al.* 2015). Homogeneous tree densities across regions likely arise from irrigation in arid regions and mowing, trimming, and thinning in more mesic regions, resulting from human preferences for savanna-like landscapes in urban regions (Balling & Falk 1982; Orians & Heerwagen 1992; Falk & Balling 2009).

We found evidence that one of the measured species’ functional traits did respond to the aridity gradient. In particular, the ratio of leaf perimeter to surface area, which has been empirically associated with leaf hydraulic traits (Sack *et al.* 2003), increased with increasing aridity. Larger values indicate either more lobed or smaller leaves, both of which decrease boundary layer resistance and allow leaves to cool more quickly in hot environments (Givnish & Vermeij 1976). Critically, we found no evidence for systematic variation in how this trait responded to the environment across different vegetation pools: the cultivated and natural pools responded equally to the aridity gradient, perhaps because the problem of excess heat load on leaves is not alleviated by irrigation, and the cooling benefit from leaves with high values is relevant to all species pools. Thus our results are not solely consistent with a homogenisation of traits (*e.g.*, Cadotte *et al.* 2009; Peppe *et al.* 2011). We found no evidence for differences in the variance of any structural (or diversity) metrics within urban and natural assemblages: variation among the urban pools was comparable to variation among the natural pools.

### Broader Implications

We show that evaluating whether urban plant floras are homogenised requires explicit consideration of both the vegetation pool (cultivated, spontaneous) relative to natural reference areas, as well as the attribute of the pool being evaluated (diversity, composition, structure). Across regions, urban species pools resemble each other more strongly than do natural area pools in terms of species and phylogenetic composition, providing evidence for homogenisation. Yet species within the natural pools were found within urban pools, implying exchange of species between pools that results in shared species and lineages. Notably, spontaneous pools are intermediate in composition between cultivated and natural pools (Figure 3), indicating that they may operate as an exchange reservoir that serves as a sink for cultivated species and as both a source and a sink for species in natural areas. Comparing urban ecosystems with natural ecosystems allows us to tease apart different dimensions of urban biodiversity, unpacking the influence of human desires (for trees) and environmental drivers (for thinner leaves). If, as we have argued, the natural and spontaneous pools have species in common, there is hope that urban areas can act as reservoirs of biodiversity and thus save species from extinction. At the same time, natural areas will increasingly receive species from urban pools, shifting the composition and diversity of the regional and continental floras to reflect the vegetation preferred and readily cultivated by humans. If composition within urban pools is comparable to natural areas, perhaps cities can reflect the eco-regions within which they are embedded. Whether the novel ecosystems that result from the merger of natural and cultivated species will permit stable, productive assemblages, however, remains to be seen.

